# Tyramine receptor drives olfactory response to (*E*)-2-decenal in the stink bug *Halyomorpha halys*

**DOI:** 10.1101/2020.10.05.326645

**Authors:** Luca Finetti, Marco Pezzi, Stefano Civolani, Girolamo Calò, Chiara Scapoli, Giovanni Bernacchia

**Affiliations:** Department of Life Sciences and Biotechnology, University of Ferrara, Ferrara, Italy; InnovaRicerca s.r.l. Monestirolo, Ferrara, Italy; Department of Biomedical and Specialty Surgical Sciences, Section of Pharmacology, University of Ferrara, Ferrara, Italy

**Keywords:** Brown Marmorated Stink Bug, TAR1 receptor, Antennae, Olfaction, Behavior, RNAi

## Abstract

In insects, the tyramine receptor 1 (TAR1) has been shown to control several physiological functions, including olfaction. We investigated the molecular and functional profile of the *Halyomorpha halys* type 1 tyramine receptor gene (HhTAR1) and its role in olfactory functions of this pest. Molecular and pharmacological analyses confirmed that the *HhTAR1* gene codes for a true TAR1. The RT-qPCR analysis revealed that *HhTAR1* is expressed mostly in adult brain and antennae as well as in early development stages (eggs, 1^st^ and 2^nd^ instar nymphs). In particular, among the antennomeres that compose a typical *H. halys* antenna, *HhTAR1* was more expressed in flagellomeres. Scanning electron microscopy (SEM) investigation revealed the type and distribution of sensilla on adult *H. halys* antennae: both flagellomeres appear rich in trichoid and grooved sensilla, known to be associated with olfactory functions. Through a RNAi approach, topically delivered *HhTAR1* dsRNA induced a 50 % gene downregulation after 24 h in *H. halys* 2^nd^ instar nymphs. An innovative behavioral assay revealed that *HhTAR1* RNAi-silenced 2^nd^ instar nymphs were less susceptible to the alarm pheromone component (*E*)-2 decenal as compared to control. These results provide critical information concerning the TAR1 role in olfaction regulation, especially alarm pheromone reception, in *H. halys*. Furthermore, considering the emerging role of TAR1 as target of biopesticides, this work paves the way for further investigation on innovative methods for controlling *H. halys*.

## Introduction

Identifying volatile compounds through the olfactory system allows insects to find food sources, avoid predators as well as localize putative partners and oviposition habitats (Gadenne et al., 2016). Furthermore, the olfactory modulation by volatile molecules with repellent activity could be a promising strategy for pest control (Carey & Carlson, 2011). The basic organization of the olfactory system begins with the antennae, organs possessing cuticular structures, the sensilla, innervated by olfactory sensory neurons (OSNs) (Amin & Lin, 2019). The OSNs recognize different molecules through special olfactory receptors. Each OSN expresses only one type of olfactory receptor, ensuring the specificity of signal for a single odour (Zhao & McBride, 2020). When an OSN is activated, it sends the output signal through the axon to the antennal lobe. Here, excitatory projection neurons (PNs) transport the olfactory information to brain centres such as the mushroom body and the lateral horn (Tanaka et al., 2012). The mushroom body plays an important role in the olfactory learning and memory (Caron et al., 2013) while the lateral horn controls innate olfactory response functions (Jefferis et al., 2007). In insects, the olfactory system can be modulated by exogenous (photoperiod, temperature) and endogenous (hormones) factors.

The biogenic amines tyramine (TA) and octopamine (OA) are important neurohormones and neurotransmitters that play a key role in the regulation of primary mechanisms in invertebrates such as locomotion, learning memory and olfaction (Roeder, 2005). Initially, TA was considered only as a biosynthetic intermediate of OA (Lange, 2009), but later numerous studies showed that TA is indeed an important neurotransmitter (Blenau & Baumann, 2003; Roeder, 2005; Lange, 2009; Roeder, 2020). Among invertebrates, TA is the endogenous agonist of the tyramine receptors (TARs). Structurally, TARs receptors are part of the superfamily of G protein-coupled receptors sharing the typical structure with seven transmembrane domains (Ohta & Ozoe, 2014). Several studies have highlighted that TARs can by coupled with both G_q_ (increasing intracellular calcium levels) and G_i_ proteins (decreasing cAMP levels) (Saudou et al., 1990; Blenau et al., 2000; Enan, 2005; Rotte et al., 2009). Based on the rank order of potency of agonists, the TAR receptors have been classified into three different types (Wu et al., 2014): TAR1 and TAR2, coupled with Gq and Gi and proteins, while TAR3 has been so far described only in *Drosophila melanogaster* (Bayliss et al., 2013; Wu et al., 2014). The first TAR1 was characterized in 1990 in *D. melanogaster* (Saudou et al., 1990). The receptor, called Tyr-dro, showed higher affinity (12-fold) for TA than for OA and was mainly expressed in heads. Since then the same receptor has been characterized in several orders of insects: Hymenoptera (Blenau et al., 2000), Orthoptera (Poels et al., 2001), Lepidoptera (Ohta et al., 2003), Hemiptera (Hana & Lange, 2017a) and Diptera (Finetti et al., 2020). Several physiological and behavioral functions are controlled by TAR1, including olfaction. In 2000 Kutsukake et al. characterized *honoka*, a *D. melanogaster* strain that presented a TAR1 mutation and a compromised olfactory profile. These insects were not able to localize repellent stimuli suggesting that TAR1 could be involved in this physiological response. Furthermore, RNAi-mediated modulation of TAR1 expression was shown to affect the gregarious and solitary phase change through a different olfactory sensibility to attractive and repulsive volatiles (Ma et al., 2015). In honeybee antennae, an upregulation of TAR1 was observed during the transition from nurses to pollen foragers, suggesting a TAR1-regulation in their behavioral plasticity (McQuillan et al., 2012). High TAR1 levels were also found in the antennae of *Mamestra brassicae* and *Agrotis ipsilon*, further suggesting a pivotal role of this receptor in olfactory modulation (Brigaud et al., 2009; Duportets et al., 2010). The TAR1s are considered interesting target for insecticides, especially bioinsecticides. Amitraz is an acaricide and non-systemic insecticide that targets the OA receptors. However, recent studies have shown that Amitraz can exert its toxic effect also through TAR1 activation (Wu et al., 2014; Kumar, 2019). Furthermore, a secondary metabolite of Amitraz, BTS-27271, increases the TA response on the *Rhipicephalus microplus* TAR1 (Gross et al., 2015). Concerning biopesticides, in the last years several studies have demonstrated that monoterpenes directly activate TAR1. In particular, Enan (2005) was the first to describe an agonist effect of several monoterpenes (thymol, carvacrol, α-terpineol, eugenol) on the *D. melanogaster* TAR1. However, the same monoterpenes did not show the same pharmacological profile on *D. suzukii* and *R. microplus* TAR1 receptors where they act as positive allosteric modulators (Gross et al., 2017; Finetti et al., 2020).

*H. halys* (Rhyncota; Pentatomidae) is an insect typical of the Eastern Asia (China, Japan, Taiwan, and Korea) (Haye et al., 2015), was detected for the first time in USA in 1998 (Hoebeke & Carter, 2003) and became a stable presence in orchards since 2010 (Rice et al., 2014). Its first European appearance was reported in 2004 in Switzerland then leading to its spread across the continent (Cesari et al., 2018). *H. halys* is responsible for major damages to many economically relevant crops (Leskey & Nielsen, 2018). The damages are caused by the perforation of the external integuments of fruits by the rostrum, the specialized sucking apparatus typical of Rhynchota. This causes necrotic areas on fruits, as well as the transmission of other phytopathogens, leading to a relevant devaluation of the product (Peiffer & Felton, 2014). In the Asiatic regions, the life cycle of *H. halys* consists of only one generation per year (Lee et al., 2013). However, in warmer regions, the insect is able to complete up to four annual generations, significantly increasing its number in the area (each female is able to lay between 100 and 500 eggs for cycle) (Nielsen et al., 2016). This particular phytopathogen shows high resistance to common pesticides, making difficult its control and elimination (Bergmann & Raupp, 2014).

The present work describes the role of TAR1 in *H. halys* olfaction. Through a RNAi-mediated knockdown of *HhTAR1* expression, the olfaction response to the alarm pheromone component (*E*)-2-decenal was studied with an innovative behavioral assay. These findings shed light on the importance of the TAR1 receptor in *H. halys* and might contribute to develop new control tools against this pest.

## Materials and Methods

### Insects and Reagents

Individuals of *H. halys* were reared on green beans and kiwi with a photoperiod of 16 h light: 8 h dark, at a temperature of 24 ± 1 °C. Tyramine hydrochloride, octopamine hydrochloride, yohimbine hydrochloride, γ-aminobutyric acid, serotonin hydrochloride, epinephrine, norepinephrine, brilliant black, Bovine Serum Albumin (BSA), probenecid, 4-(2-hydroxyethyl)-1-piperazineethanesulfonic acid (HEPES), (*E*)-2-decenal were all obtained from Sigma-Aldrich (St. Louis, Missouri, USA). Dopamine was obtained from Tocris Bioscience (Bristol, United Kingdom). Pluronic acid and fluorescent dye Fluo-4 AM were purchased from Thermo Fisher Scientific (Waltham, Massachusetts, USA). All compounds were dissolved in dimethyl sulfoxide (10 mM) and stock solutions were kept at -20 °C until use. Serial solutions were made in the assay buffer (Hanks’ Balanced Salt solution (HBSS)/HEPES 20 mM buffer, containing 0.01 % BSA and 0.1 % DMSO).

### Isolation and cloning of full-length HhTAR1

Total RNA was extracted from four adults of *H. Halys* using RNAgent® Denaturing Solution (Promega, Madison, Wisconsin, USA), quantified in a micro-volume spectrophotometer Biospec-Nano (Shimadzu, Kyoto Japan) and analysed by 0.8 % w/v agarose gel electrophoresis. One µg of RNA was treated with DNase I (Thermo Fisher Scientific) and used for the synthesis of cDNA, carried out with the OneScript® Plus cDNA Synthesis Kit (ABM, Richmond, Canada). For amplification of the full *HhTAR1* open reading frame (ORF), specific primers were designed based on the annotated transcript (XM_014422850.2). The Kozak translation initiation sequence (GCCACC) was inserted at 5’ end of the receptor (**Table 1**). High fidelity amplification was achieved using Herculase II Fusion DNA Polymerase (Agilent, Santa Clara, California, USA) and a touchdown thermal profile: predenaturation at 95 °C for 3 mins, followed by 10 cycles at 95 °C for 20 s, 65-55 °C for 20 s (minus 1 °C/cycle), 68 °C for 2 mins, 30 cycles at 95 °C for 20 s, 55 °C for 20 s, 68 °C for 2 mins and a final extension at 68 °C for 4 mins. The PCR product was gel purified by Illustra GFX PCR DNA and Gel Band Purification Kits (GE Healthcare, Chicago, Illinois, USA), cloned into pJET 1.2/blunt vector (Thermo Fisher Scientific) and transformed into *E*.*coli* SIG10 5-α Chemically Competent Cells (Sigma-Aldrich). Positive clones were selected using LB broth agar plates with 100 µg/ml ampicillin. Plasmid was then extracted by GenElute™ Plasmid Miniprep Kit (Sigma-Aldrich) and verified by DNA sequencing (BMR Genomics, Padua, Italy). The sequence, named *HhTAR1*, was deposited in GenBank with the accession number MT513133. For expression in Human Embryonic Kidney (HEK 293) cells, the open reading frame of *HhTAR1* was excised from pJET 1.2 vector and inserted into the pcDNA 3.1 (+) Hygro expression vector using *Xho I* and *Xba I* restriction sites.

**Table 1.**
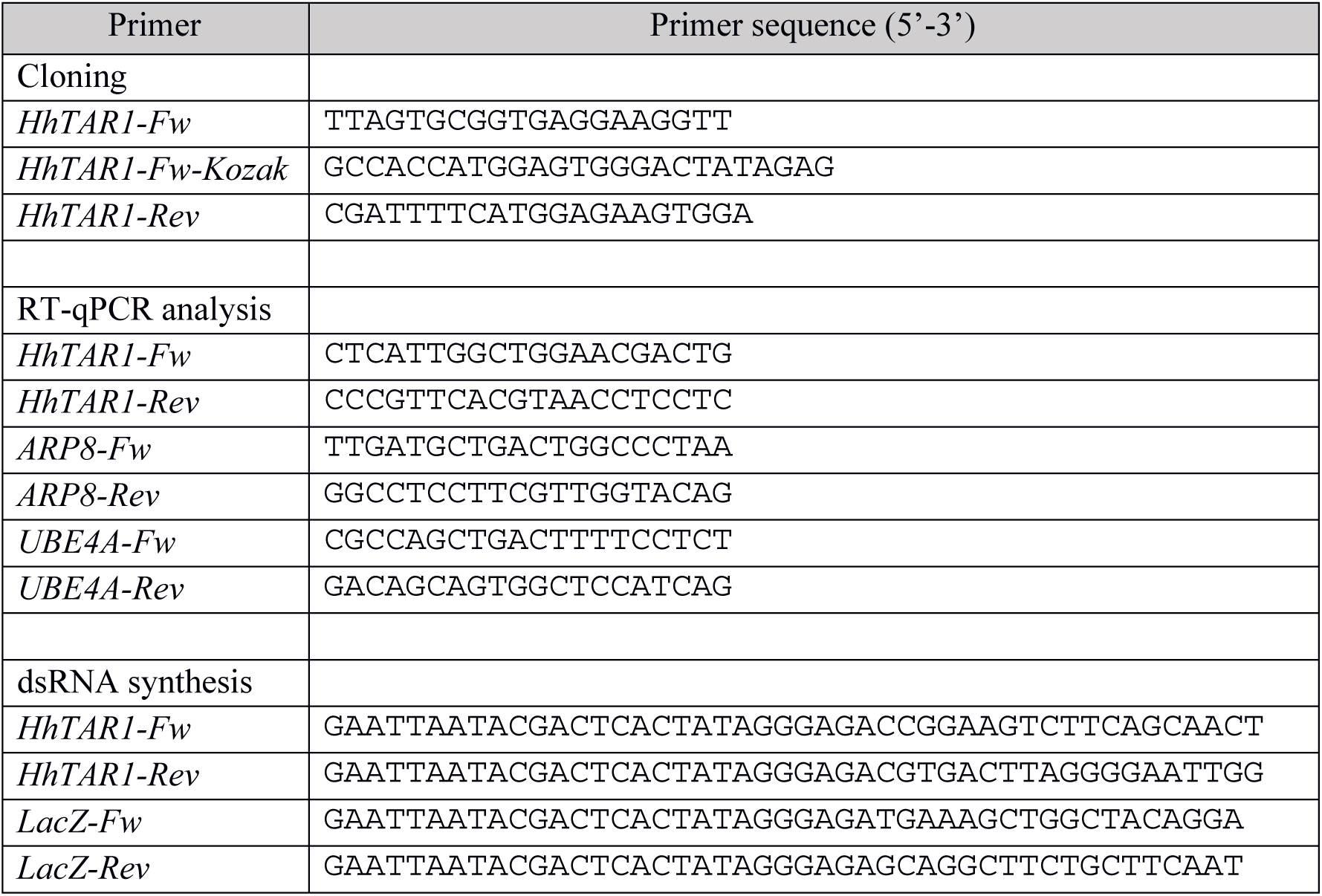
Primers used in this study.

### Multiple sequence alignment and general bioinformatics analysis

Multiple protein sequence alignments between the deduced amino acid sequence of HhTAR1 and other type 1 tyramine receptor sequences were performed using Clustal Omega (https://www.ebi.ac.uk/Tools/msa/clustalo/) and BioEdit Sequence Alignment Editor 7.2.6.1. Phylogenetic neighbour-joining analysis was performed by MEGA software (version 7) with 1000-fold bootstrap resampling. The *D. melanogaster* GABA B receptor (GABABR) was used as an outgroup to root the tree.

### HhTAR1 transient expression in HEK 293

HEK 293 cells were grown at 37 °C and 5 % CO_2_ in Dulbecco’s modified Eagles medium high glucose (D-MEM) supplemented with 10 % fetal bovine serum (Euroclone, Milan, Italy). To prevent bacterial contamination, penicillin (100 U/ml) and streptomycin (0.1 mg/ml) were added to the medium. The cells were transiently transfected with pcDNA 3.1 (+) / HhTAR1 in T75 cell culture flasks (Euroclone) using JetOPTIMUS (Polyplus-Transfection, New York, New York, USA), following the manufacturer’s protocol. Cells were incubated in the transfection medium for 24 h at normal cell growth conditions before their use for the calcium mobilization assay.

### Calcium Mobilization Assay

Cells were seeded at a density of 50,000 cells per well, total volume of 100 µl, into poly-D-lysine coated 96-well black, clear-bottom plates. After 24 h incubation at normal cell culture condition, the cells were incubated with HBSS 1X supplemented with 2.5 mM probenecid, 3 μM of the calcium sensitive fluorescent dye Fluo-4 AM and 0.01 % pluronic acid, for 30 mins at 37 °C. After that, the loading solution was removed and HBSS 1X supplemented with 20 mM HEPES, 2.5 mM probenecid and 500 μM brilliant black were added. Cell culture and drug plates were placed into the fluorometric imaging plate reader FlexStation II (Molecular Devices, Sunnyvale, California, USA) and fluorescence changes were measured after 10 mins of stabilization at 37 °C. On-line additions were carried out in a volume of 50 μl/well after 20 s of basal fluorescence monitoring. In antagonism protocols, to facilitate drug diffusion into the wells the assays were performed at 37 °C with three cycles of mixing (25 μl from each well moved up and down three times). The fluorescence readings were measured every two s for 120 s.

### Quantitative real-time PCR analysis

Total RNA was extracted from *H. halys* samples at various developmental stages (eggs, 1^st^ to 5^th^ instar nymphs, adult males and females) and different organs (antennae, brain, midgut, reproductive organs) using RNAgent® Denaturing Solution (Promega). The organs of *H. halys* were dissected in a RNA preservation medium (20 mM EDTA disodium (pH 8.0), 25 mM sodium citrate, 700 g/l ammonium sulphate, final pH 5.2). One µg of purified RNA was then treated with DNase I (Thermo Fisher Scientific) and used for cDNA synthesis, carried out with OneScript® Plus cDNA Synthesis Kit (ABM), according to the manufacturer’s instructions. Real time PCR was performed using a CFX Connect Real-Time PCR Detection System (Bio-Rad) in a 12 µl reaction mixture containing 0.8 µl of the cDNA obtained from 1 µg of total RNA, 6 µl ChamQ SYBR qPCR Master Mix (Vazyme, Nanchino, China), 0.4 µl forward primer (10 µM), 0.4 µl reverse primer (10 µM) and 3.6 µl nuclease free water. Thermal cycling conditions were: 95 °C for 2 mins, 40 cycles at 95 °C for 15 s and 60 °C for 20 s. After the cycling protocol, a melting-curve analysis from 60 °C to 95 °C was applied. Expression of *HhTAR1* was normalized in accordance with the relative quantitation method (Larionov et al., 2005) using *ARP8* and *UBE4A* as reference genes (Bansal et al., 2016). Gene-specific primers (**Table 1**) were used and at least three independent biological replicates, made in triplicate, were performed for each sample.

### Antennae preparation and SEM analysis

Preliminary morphological investigations were performed on ten adults of *H. halys* (five males and five females) using a Nikon SMZ 800 stereomicroscope (Nikon Instruments Europe, Amsterdam, The Netherlands), provided with a Nikon Digital Sight Ds-Fil camera (Nikon Instruments Europe, Amsterdam, The Netherlands) and connected to a personal computer with the imaging software NIS Elements Documentation (Nikon Instruments Europe, Amsterdam, The Netherlands). Based on stereomicroscope observations, the head was dissected from body and prepared for scanning electron microscopy (SEM), according to previously published procedures (Pezzi et al. 2015, 2016). Afterwards, samples were critical point dried in a Balzers CPD 030 dryer (Leica Microsystems, Wetzlar, Germany), glued on stubs and coated with gold-palladium in an S150 Edwards sputter coater (HHV Ltd, Crawley, United Kingdom). The SEM observations were conducted at the Electronic Microscopy Centre of the University of Ferrara, using a Zeiss EVO 40 SEM (Zeiss, Milan, Italy).

### Synthesis of dsRNA and *H. halys* treatment

For RNAi silencing, *HhTAR1* and *LacZ* (control) amplicons, 400-500 bp long, were generated by PCR using primers with 5’ extensions containing T7 promoters (**Table 1**). These products were cloned into pJET 1.2 vector (Thermo Fisher Scientific) and then used as templates for *in vitro* dsRNA synthesis performed by T7 RNA Polymerase (Jena Bioscience, Jena, Germany), according to the manufacturer’s protocol. After one hour of synthesis at 37 °C, a DNase I (Thermo Fisher Scientific) treatment was performed and the dsRNA was clean up by ammonium acetate precipitation (Rouhana et al., 2013). Finally, the dsRNA was resuspended in ultrapure water and quantified by Biospec-Nano spectrophotometer. To induce RNAi silencing 2^nd^ stage nymphs of *H. halys* 3 days post-ecdysis were treated with 100 ng of dsTAR1 or dsLacZ in 1 µl of solution using a 0.1-2 µl micropipette. The dsRNA molecules were topically delivered through a drop placed on the abdomen of nymphs (**Supplementary figure 4**). Insects were tested by behavioral assay after 24 h while the *HhTAR1* transcript level was measured by RT-qPCR, as described above.

### Repellence assay

An open petri dish (90 mm x 15 mm), containing 24 h starved *H. halys* 2^nd^ instar nymphs and a green bean, was placed inside a plexiglas box (50 cm each side) with two lateral openings covered by nets to allow air circulation. The negative control acetone or the positive repellent control (*E*)-2-decenal were applied to a filter paper (1 cm x 1 cm) that was placed under the green bean. The positive control (*E*)-2-decenal, dissolved in acetone, was tested at a fixed quantity of 10 µg, a value ensuring the maximum repellence activity against the *H. halys* nymphs (Zhong et al., 2018). The number of *H. halys* nymphs standing and feeding on the green bean was monitored every ten minutes for one hour. Four biological replicates were made, each comprising at least ten insects, for both untreated and dsRNA treated *H. halys* nymphs. All experiments were performed in the morning in a behavioral room with a controlled temperature of 24 ± 1 °C.

### Data analysis and terminology

All data were elaborated using Graph Pad Prism 6.0 (La Jolla, California, USA). Data are expressed as mean ± SEM of n experiments and were analysed using one- or two-way analysis of variance (ANOVA) followed by Dunnett’s or Turkey’s test for multiple comparison. In the pharmacological assays, the concentration-response curves were fitted using the four parameters log logistic equation:

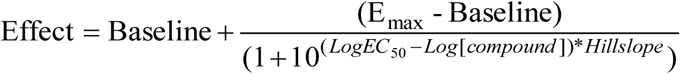

Agonist potency was expressed as pEC_50_, defined as the negative logarithm to base 10 of the agonist molar concentration that produces 50% of the maximum possible effect of that agonist. Antagonist potency was derived from Gaddum Schild equation:

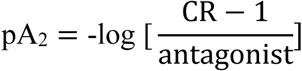

Assuming a slope value equal to unity, where CR indicates the ratio between agonist potency in the presence and absence of antagonist (Kenakin, 2014).

## Results

### Molecular characterization of HhTAR1

The amplified *HhTAR1* sequence was 1347 bp long and coded for a 449 aa polypeptide with a predicted MW of 50.97 KDa and pI of 9.41. About structural domains, both TMHMM v 2.0 software and the Kyte and Doolittle method (Kyte & Doolittle, 1982) suggest seven putative transmembrane domains, as expected for a GPCR. The helixes are flanked by an extracellular N-terminus of 51 residues and an intracellular C-terminus of 18 residues. Furthermore, the HhTAR1 sequence contains a DRY conserved sequence in the TM3, several N-glycosylation sites in the extracellular N-terminus and P-glycosylation sites, 2 specific for PKA and 10 specific for PKC (**Supplementary Figure 1**). These features are important for the correct folding and function of GPCRs (Nørskov-Lauritsen & Bräuner-Osborne, 2015). Moreover, at position 128 in TM3 there is a conserved aspartic acid (D^128^, shown in **Figure 1, panels A and B**) responsible for the interaction with TA, the endogenous agonist of the TAR1s (Ohta & Ozoe, 2014).

**Figure 1.**
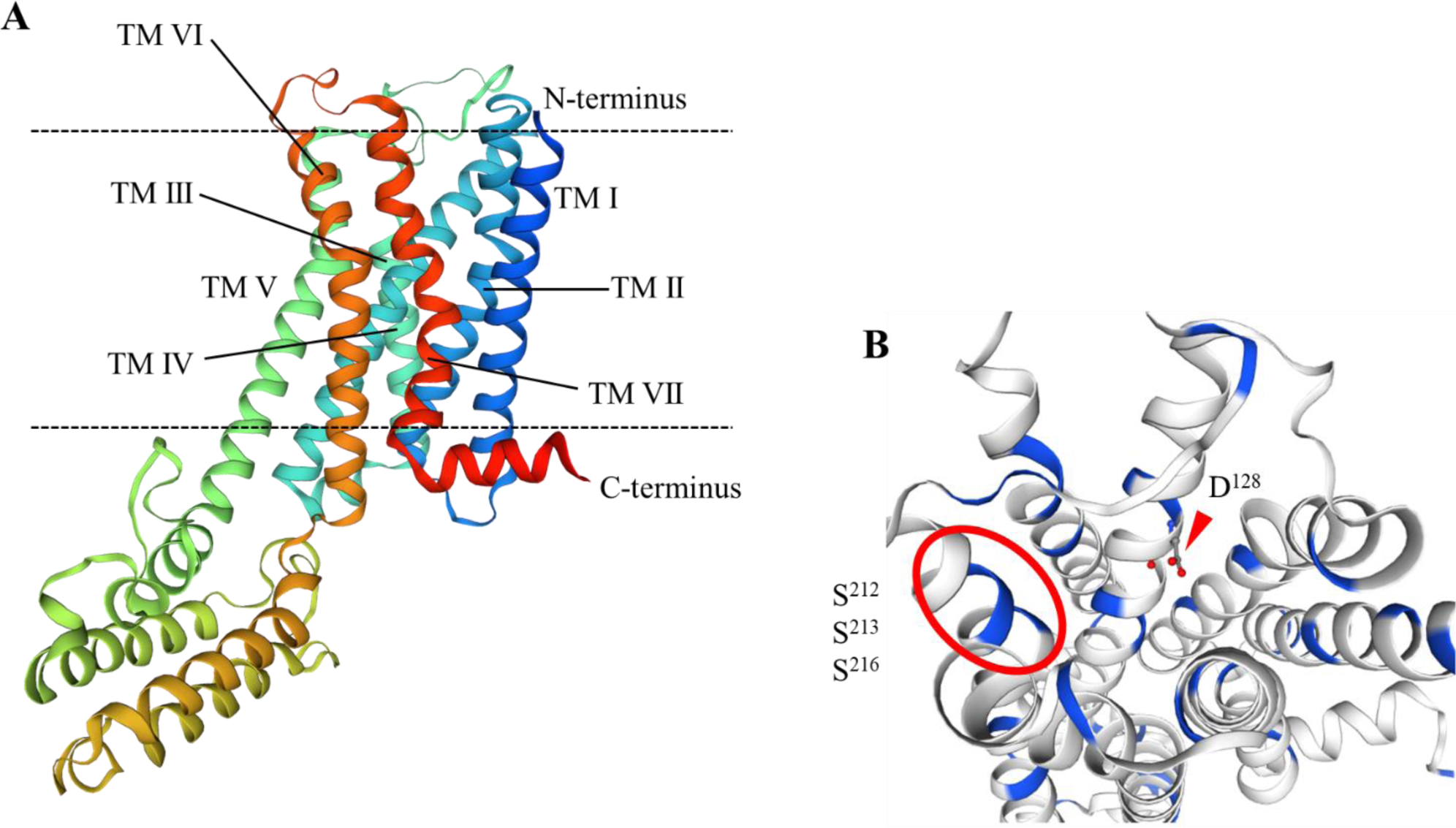
Structural overview of HhTAR1 predicted by SWISS-MODEL. (**A**) Model of the whole receptor showing the transmembrane domains. (**B**) Detail of the putative ligand binding pocket, seen from the extracellular side, of HhTAR1. All serine residues are highlighted in blue. The aspartic acid in TM3 (D^128^) is shown by a triangle and the three serine residues interacting with TA are highlighted by a circle.

To study the binding site structure, the HhTAR1 aminoacidic sequence was analysed by SWISS-MODEL (Waterhouse et al., 2018). The model was created based on the crystal structure of the human α2A adrenergic receptor (Template code: 6kux.1.A) that shares 33.51 % of sequence identity with HhTAR1. The three-dimensional model of the whole receptor and the putative ligand binding pocket are shown in **Figure 1**. In 2004, several serine residues in TM V were found to play a key role in stabilizing the interaction with TA in *Bombyx mori* and *Sitophilus oryzae* (Ohta et al., 2004; Braza et al., 2019) TAR1. These serine residues localized in the TM V are also conserved in HhTAR1 at positions S^212^, S^213^ and S^216^ (**Figure 1, panel B**). MolProbity model quality investigation (**Table 2**) confirmed the validity of the SWISS-MODEL 3D model of HhTAR1 (Chenn et al., 2010). The HhTAR1 deduced amino acid sequence was then compared to several OA and TA receptors allowing the construction of a neighbour-joining phylogenetic tree by MEGA 7 server (**Supplementary Figure 2**). As expected, HhTAR1 grouped in the TAR1 family, the main monophyletic group, and shared the highest percentage of identity with the *Rhodnius prolixus* TAR1 (Accession number: MF377527.1), another Pentatomidae. Based on the phylogenetic results, a multiple sequence alignment was performed between the HhTAR1 deduced amino acid sequence and TAR1 from other insects (**Figure 2**). The analysis further strengthen the similarity of HhTAR1 with known TAR1 receptors showing the typical GPCR structure with highly conserved domains corresponding to the transmembrane regions as well as the TA binding site.

**Table 2.** MolProbity results based on the HhTAR1 3D model obtained by SWISS-MODEL software.

| MolProbity Parameter | Result |
| --- | --- |
| MolProbity Score | 1.95 |
| ClashScore | 3.07 (M<sup>260</sup>, K<sup>263</sup>) |
| Ramachandran Favoured | 97.94 % (goal: > 98 %) |
| Ramachandran Outliers | 0.51 % (D<sup>331</sup>, P<sup>77</sup>) (goal: < 0.2 %) |

**Figure 2.**
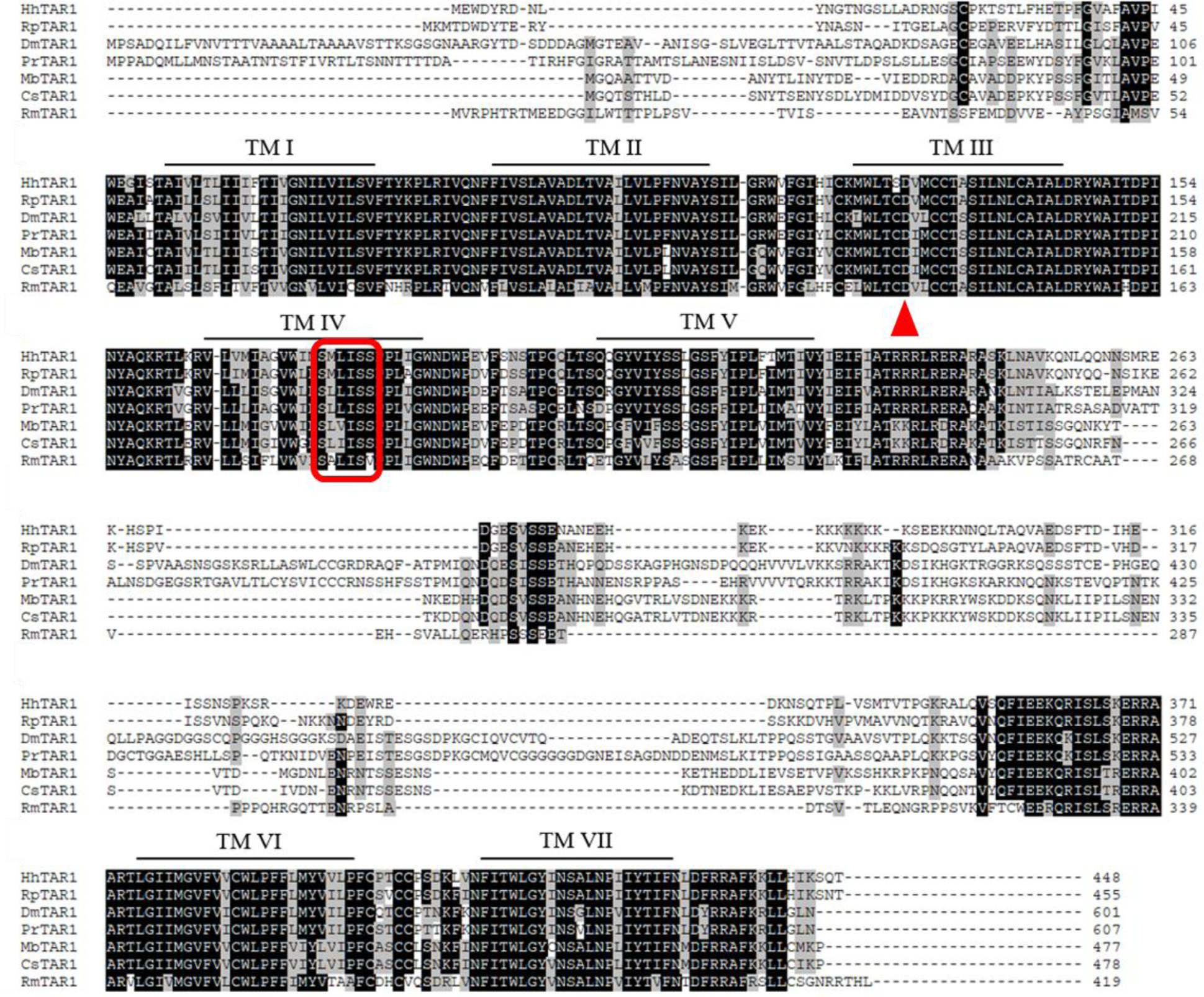
Amino acid sequence alignment of HhTAR1 with orthologous receptors from *R. prolixus* (RpTAR1), *D. melanogaster* (DmTAR1) *Phormia regina* (PrTAR1), *Mamestra brassicae* (MbTAR1), *Chilo suppressalis* (CsTAR1) and *Rhipicephalus microplus* (RmTAR1). The putative seven transmembrane domains (TM I-VII) are indicated with a black line. Identical residues are highlighted in black while conservative substitutions are shaded in grey. A red triangle indicates the conserved aspartic acid D^128^ and the serine residues that could interact with TA are shown by a red box.

### HhTAR1: pharmacological validation

In the calcium mobilization assay HhTAR1 was activated by both TA and OA in a concentration-dependent manner (**Figure 3, panel A**). TA evoked the release of intracellular calcium with pEC_50_ values of 5.99 (CL_95%_ 5.32-6.66) and E_max_ of 109.33 ± 14.86, while OA resulted less potent with a pEC_50_ of 4.41 (4.17-4.64) calculated assuming the TA maximum effect (**Figure 3, panel A**). In wild type HEK 293 cells, TA and OA were completely inactive when tested in the same concentration range (from 10^−10^ M to 10^−4^ M) (data not shown). Yohimbine was completely inactive as agonist, while, at 1 μM yohimbine, elicited a rightward shift of the concentration response curve to TA (**Figure 3, panel B**); a pA_2_ of 8.26 was calculated from these experiments.

**Figure 3.**
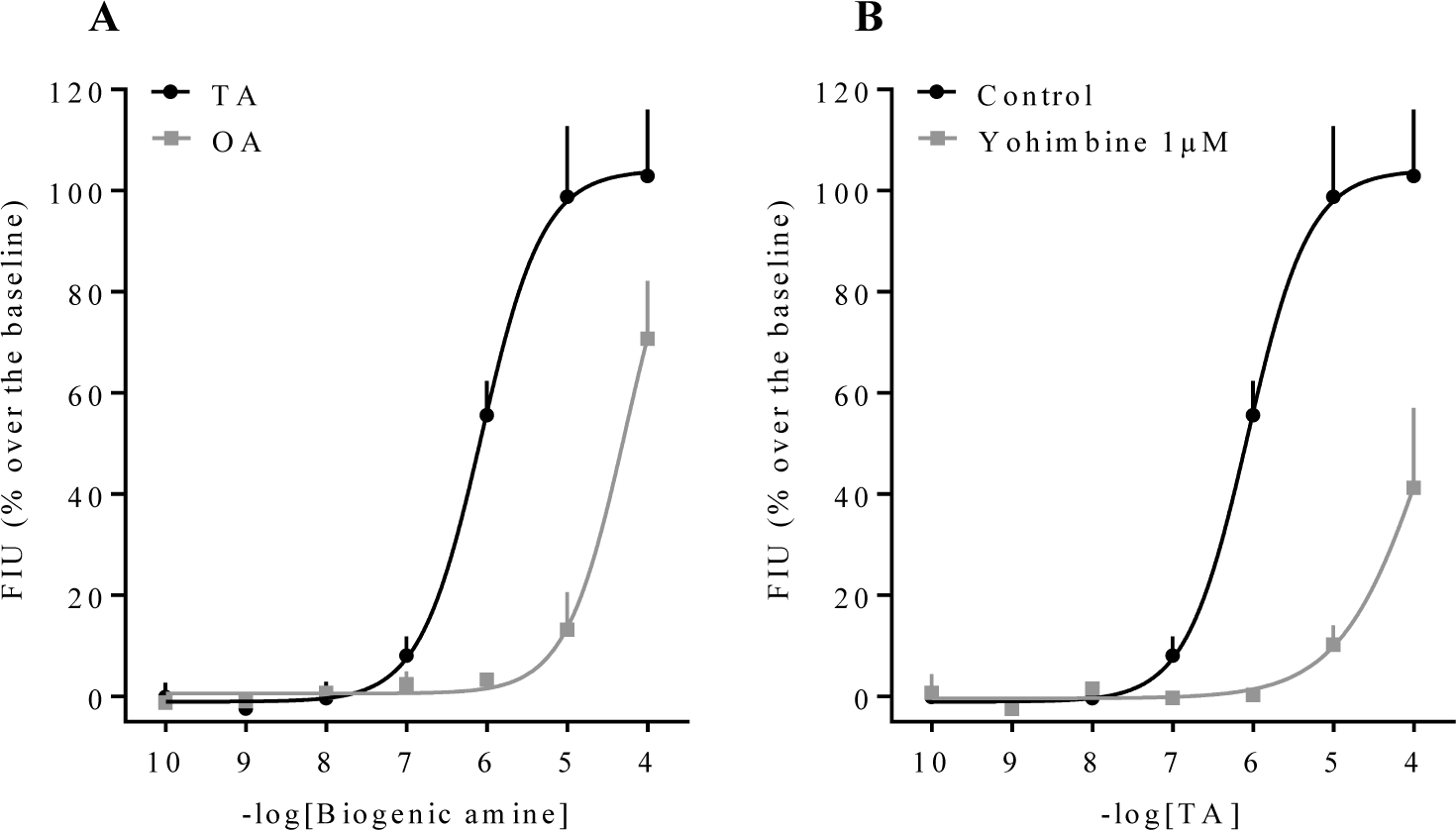
Calcium mobilization assay in HhTAR1-transfected HEK293 cells. Concentration-response curves to TA and OA (**A**). Concentration-response curves to TA in the absence (control) and in presence of 1 μM yohimbine (**B**). Data are means ± SEM of three separate experiments performed in duplicate.

In order to confirm the HhTAR1 sensibility to TA and OA, other biogenic amines such as dopamine, L-DOPA, epinephrine, norepinephrine and serotonin or important neurotransmitter like γ-aminobutyric acid were tested at 10^−4^ M as putative ligands. TA and OA were able to generate a potent effect against the receptor while the other molecules did not elicit any release of calcium (**Supplementary Figure 3**).

### *HhTAR1* expression pattern

Given the importance of TAR1s in insect physiology and behavior, *HhTAR1* expression profile was studied in all *H. halys* development stages (egg, 1^st^ to 5^th^ instar nymphs, L1 to L5, and adult) as well as in the major organs of the adult. The analysis revealed that *HhTAR1* was mostly expressed in eggs and in 1^st^ and 2^nd^ instar nymphs, with a dramatic decrease in receptor mRNA levels in the later stages from the 3^rd^ instar nymph to adult (**Figure 4, panel A**). This mRNA reduction in 2^nd^ and 3^rd^ instar nymphs was further investigated. The nymphs were divided in two parts: head + antennae and thorax + abdomen and the *HhTAR1* expression levels analysed. The *HhTAR1* mRNA level decrease affected both sections of 2^nd^ and 3^rd^ instar nymphs with different intensity (**Figure 4, panel B**): the level in head/antennae decreased only by 38% between L2/L3, while it dropped by 82 % in thorax/abdomen. This reveals that *HhTAR1* levels remain high in the nervous tissues while they decrease significantly in the rest of the nymph body. Among the different organs analysed (antennae, brains, midguts and gonads), the highest levels of *HhTAR1* transcript were detected in the brains and the antennae of both sexes, even if they were statistically more abundant in male tissues (**Figure 4, panel C**). Furthermore, *HhTAR1* expression was investigated in all antennomeres of *H. halys*. The antenna is in fact composed by a scape (SC), two pedicels (PE1 and PE2) and two flagellomeres (FL1 and FL2) (**Figure 4, panel E**). The *HhTAR1* mRNA was detected in all antennomeres but it was 2-3 times more abundant in FL1 and FL2 in comparison to SC and both elements of pedicel (**Figure 4, panel D**).

**Figure 4.**
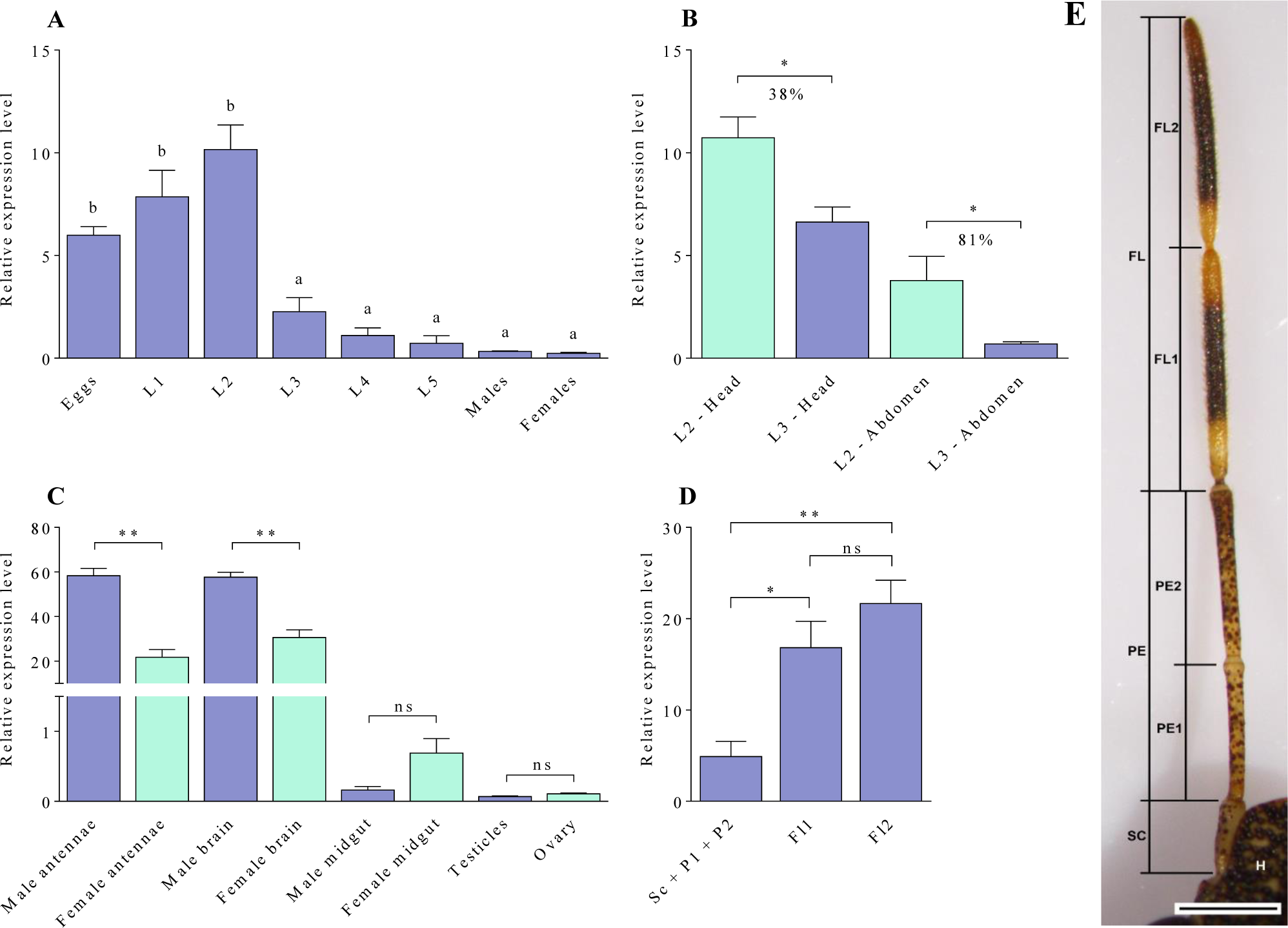
mRNA expression levels of *HhTAR1* gene. (**A**) Expression of *HhTAR1* gene in all development stages: eggs, 1^st^ to 5^th^ instar nymphs (L1 to L5), adult male and female. (**B**) Expression of *HhTAR1* in different parts of 2^nd^ (L2) and 3^rd^ (L3) *H. halys* instar nymphs. (**C**) Expression of *HhTAR1* gene in organs of both sexes. (**D**) Expression of *HhTAR1* in different parts of adult *H. halys* antennae. Data represent means ± SEM of at least three independent experiments performed in triplicate. * p < 0.05 ** p < 0.01 according to one-way ANOVA followed by multiple comparisons Bonferroni post-hoc. (**E**) Antenna structure of the adult *H. halys* observed on a stereomicroscope. FL, flagellum; FL1, first segment of flagellum; FL2, second segment of flagellum; H, head; PE, pedicel; PE1, first segment of pedicel; PE2, second segment of pedicel; SC, scape.

### Sensilla investigation by SEM

The different expression of *HhTAR1* in antennomeres required a further characterization of the antenna. The antennae, the main organs of the olfactory system in insects, are rich in sensilla whose morphology correlates with their physiological role. We investigated by scanning electron microscopy (SEM) the morphology and distribution of sensilla in the different parts of adult *H. halys* antennae: scape (SC), two pedicels (PE1 and PE2) and two flagellomeres (FL1 and FL2) (**Figure 4, panel E**). In the SC and both PEs, sporadic basiconic sensilla (BS) (**Figure 5, panels A, B and E**) were visible along with particular perforations classified as pit sensilla (PT), or coeloconic sensilla, found in both PEs (**Figure 5, panels F and G**). Several chaetic sensilla (CH) were observed in the PE2-FL1 junction area (**Figure 5, panel C**). A high number of sensilla was found in both FLs, classified as trichoid (TR) (**Figure 5, panels D and I**), basiconic (BS), or grooved sensilla (**Figure 5, panel H**).

**Figure 5.**
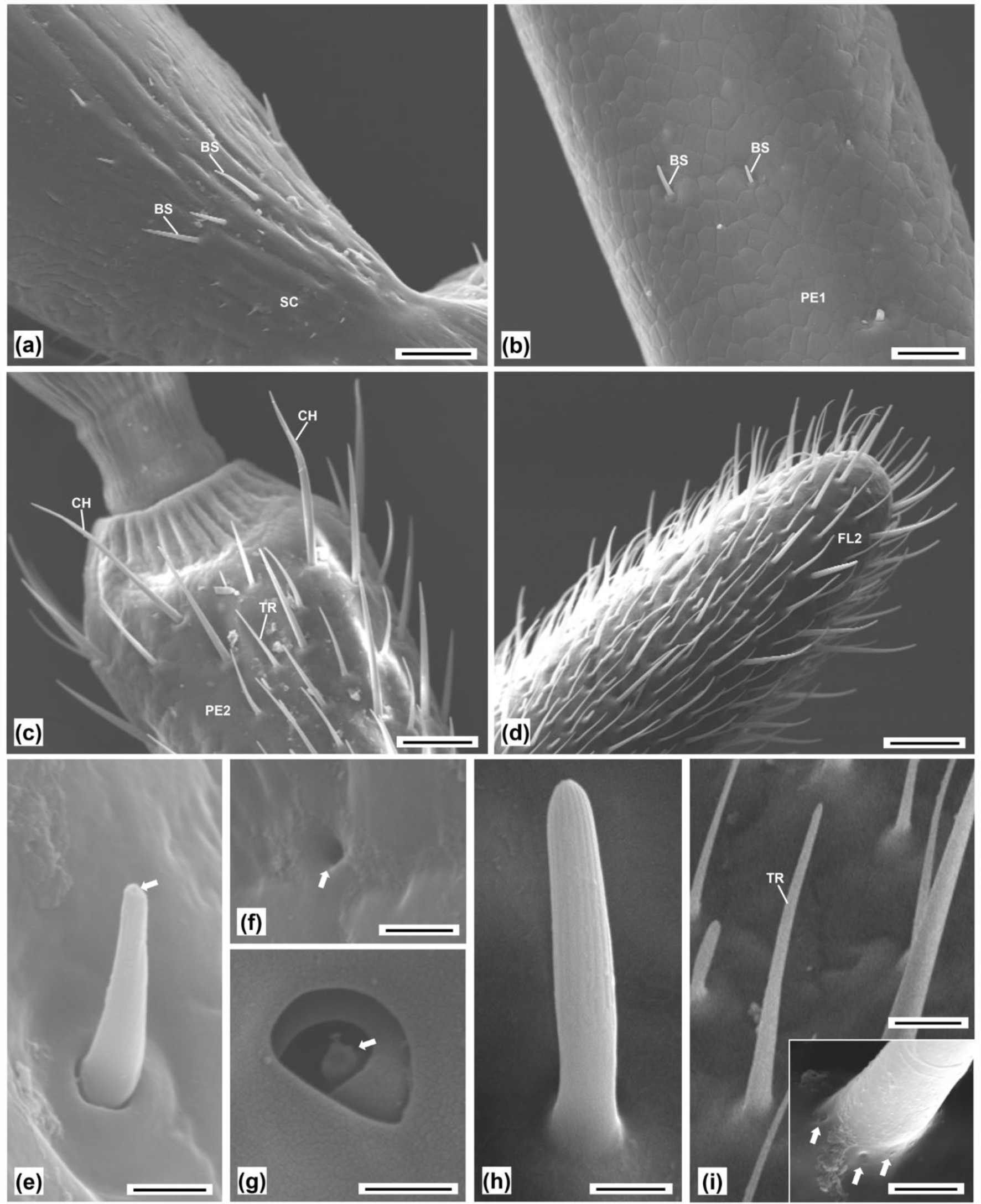
Antennae of adult *H. halys* observed at the scanning electron microscope (SEM) (**A-I**). (**A**) Female antenna, detail of the base of the scape. Scale bar 50µm. (**B**) Male antenna, detail of the first segment of the pedicel base of the scape. Scale bar 25µm. (**C**) Male antenna, distal part of the second segment of the pedicel. Scale bar 50µm. (**D**) Male antenna, tip of the second segment of the flagellum. Scale bar 50µm. (**E**) Female antenna, basiconic sensillum of the scape with a tip perforation (arrow). Scale bar 2.5µm. (**F**) Male antenna, perforation of the pedicel (arrow). Scale bar 1.5µm. (**G**) Female antenna, pit sensillum of the pedicel, containing a peg (arrow). Scale bar 1.5µm. (**H**) Female antenna, grooved sensillum of the flagellum. Scale bar 2.5µm. (**I**) Female antenna, trichoid sensillum of the flagellum. Scale bar 10µm. Inlay: detail of the base of the trichoid sensillum, showing microperforations (arrows). Scale bar 2.5µm. Abbreviations: BS, basiconic sensillum; CH, chaetic sensillum; FL2, second segment of flagellum; GR, grooved sensillum; PE1, first segment of pedicel; PE2, second segment of pedicel; SC, scape; TR, trichoid sensillum.

### *H. halys* dsRNA treatment and repellency assay

To investigate the functional role of HhTAR1 in *H. halys* behavior and chemosensory recognition firstly a behavioral repellence assay was set up. *H. halys* 2^nd^ instar nymphs were offered a green bean in the presence or absence of the alarm pheromone component (*E*)-2-decenal and the number of individuals feeding or standing on the bean was measured during a period of an hour. (*E*)-2-decenal, as expected, was able to repel approximately 50% of the nymphs compared to the acetone-treated group, used as control (**Figure 6, panel A)**. Subsequently, to assess the physiological relevance of HhTAR1 in repellency, a RNAi -silencing approach was applied to 2^nd^ instar (L2) nymphs. HhTAR1 dsRNA was administered by topical delivery (**Supplementary Figure 4**) to *H. halys* 2^nd^ instar nymphs and the silencing effect on *HhTAR1* transcript levels was evaluated by RT-qPCR 24 h after the treatment. The dsTAR1 treatment did induce a gene silencing effect, with a 50 % decrease in transcript abundance while the dsLacZ negative control RNA did not cause any variation (**Figure 6, panel B**). Interestingly, the insects treated the *HhTAR1*-dsRNA exhibited a reduced sensitivity to (*E*)-2-decenal, i.e. they moved towards and fed on green bean in the presence of (*E*)-2-decenal in a similar manner to the acetone-only control (**Figure 6, panel C**). On the other hand, the behavior of nymphs treated with *LacZ*-dsRNA was unmodified, therefore the alarm pheromone correctly repelled the insects (**Figure 6, panel D**).

**Figure 6.**
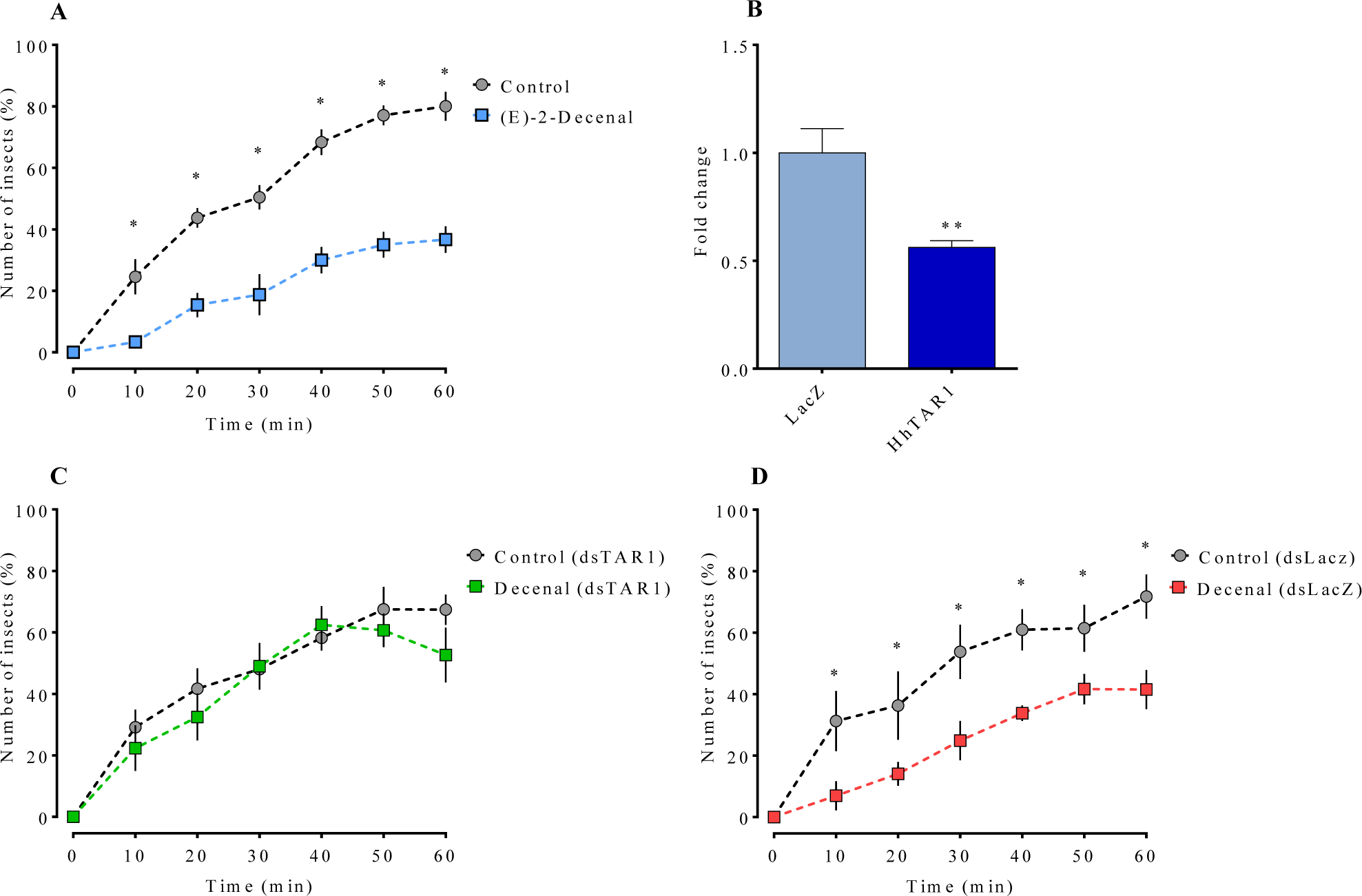
Olfactory modulation of *H. halys* 2^nd^ instar nymphs. (**A**) Behavioral repellence assay on *H. halys* 2^nd^ instar in the presence or absence of the alarm pheromone (*E*)-2-decenal. (**B**) Reduction in *HhTAR1* transcript levels by RNAi. Each bar shows the mean fold change ± SEM (standard error) of four independent replicates of *H. halys* 2^nd^ instar nymphs 24 h after gene-specific dsRNA treatment, topically delivered. LacZ specific dsRNA treatment was used as a negative control. ** p < 0.01 vs control according to student’s t test. (**C**) Behavior assay after dsHhTAR1 administration or (**D**) dsLacZ. Data are means ± SEM of four independent replicates for a total of at least 50 insect tested. * p < 0.05 vs control according to two-way ANOVA (time x treatment) followed by Dunnett post-hoc.

## Discussion

Since its appearance in Europe and in America, *Halyomorpha halys* has caused serious damage to agriculture (Rice et al., 2014; Valentin et al., 2017). Due to its reduced susceptibility to traditional control strategies, new methods for *H. halys* containment need to be developed, identifying innovative chemical compounds as well as new targets based on biochemistry, physiology and behavior of this insect.

This study deal with the molecular and pharmacological characterization of the *H. halys* type 1 tyramine receptor (HhTAR1). Through a RNAi silencing of *HhTAR1*, it was possible to reveal the important role of HhTAR1 in physiological aspects of *H. halys*, such as the olfactory response to the alarm pheromone (*E*)-2-decenal.

The HhTAR1 polypeptide shares many structural features with TAR1s from other insect (Ohta & Ozoe, 2014). HhTAR1 contains seven highly conserved transmembrane segments, as expected for a GPCR, as well as phosphorylation and glycosylation sites, typical for this receptor class and essential for the correct protein folding and receptor signalling (Nørskov-Lauritsen & Bräuner-Osborne, 2015; Alfonzo-Mèndez et al., 2017). Most of these sites (seven phosphorylation sites - T^235^ and S^246, 260, 294, 319, 321, 364^) are localized in the long intracellular loop between TM V and VI and are probably involved in receptor signalling and regulatory processes such as desensitization and internalization. Concerning the TA binding site, the main amino acid residue interacting with the endogenous agonist is an aspartic acid located in TM III and well conserved in all insect TAR1s as judged by alignment studies (Braza et al., 2019). In HhTAR1 this Asp residue is found at position 128 (D^128^). The D^128^ involvement in ligand binding has been confirmed in a mutation study performed on *Bombyx mori* TAR1 that showed that the orthologous Asp residue binds the TA-amine group with an ionic bond reinforced by H-bond (Ohta et al., 2004). The same study also showed that several serine residues in TM V stabilise the interaction between TAR1 and TA. The HhTAR1 molecular model furthermore suggests that three serine residues (found at position 212, 213 and 216 and well conserved within TAR1 insects family) might be involved in generating the receptor binding pocket (Ohta et al., 2004; Braza et al., 2019).

The structural description encouraged to proceed towards a functional characterization. The HhTAR1 coding region was cloned and expressed into HEK 293 cells and the recombinant receptor tested for its ability to respond to known TAR1 ligands. In the calcium mobilization assay TA was significantly more potent than OA, as observed for other TAR1s (Gross et al., 2015; Hana & Lange, 2017a; Finetti et al., 2020). Furthermore, the effect of TA was sensitive to the antagonist yohimbine, as observed in other orthologous TAR1s (Saudou et al., 1990; Gross et al., 2015; Hana & Lange, 2017a; Finetti et al., 2020). Other biogenic amines, such as dopamine and adrenaline, were not able to activate HhTAR1, also shown also in *R. microplus* TAR1 (Gross et al., 2015).

Many studies support the physiological role of TAR1 in processes such as locomotion (Saraswati et al., 2004; Schützler et al., 2019), metabolic control (Nishimura et al., 2005; Li et al., 2017; Roeder, 2020), reproduction (Hana & Lange, 2017a; Hana & Lange, 2017b) and olfaction (Kutsukake et al., 2000; Brigaud et al., 2009; Duportets et al., 2010; McQuillan et al., 2012; Ma et al., 2015; Zhukovskaya & Polyanovsky, 2017; Ma et al., 2019b). The *TAR1* expression patterns mirror its functional roles because the *TAR1* gene is highly expressed in the CNS, salivary glands and antennae in different insect species (Duportets et al., 2010; McQuillan et al., 2012; Wu et al., 2014; El-Kholy et al., 2015; Hana & Lange, 2017a; Ma et al., 2019a; Finetti et al., 2020). Two studies conducted in 2017 on the honeybee brain showed that *TAR1* was mainly expressed at the presynaptic sites in antennal lobe OSNs and in the mushroom bodies PNs, which are essential structures for the olfactory system in insects (Synakevitch et al., 2017; Thamm et al., 2017). Similarly, in *H. halys HhTAR1* appeared strongly expressed in brain and antennae, but was less expressed in the midgut and reproductive systems of adults. Furthermore, *HhTAR1* mRNA was more abundant in the male brain than in the female one. This sex-dependent *TAR1* expression was also detected in *D. suzukii* (Finetti et al., 2020) and *P. xylostella* (Ma et al., 2019a) suggesting that TAR1 could be involved in male specific functions such as development as well as reproduction. The high brain expression of *HhTAR1* correlates well with the abundance of TAR1 in CNS of numerous insect species (El-Kholy et al., 2015; Hana & Lange, 2017a; Finetti et al., 2020) where it regulates several sensory processes (Roeder et al., 2003; Lange, 2009; Ohta & Ozoe, 2014; Neckameyer & Leal, 2017). Interestingly, *HhTAR1* was also highly abundant in the antennae. As a matter of fact, several studies have shown that *TAR1* is expressed in these structures even if its role in the antennae is still unclear. A possible correlation between TAR1 and olfaction was established for the first time in 2000 (Kutsukake et al., 2000). This study characterized a *D. melanogaster* TAR1-mutant line, called *honoka*, whose behavioral responses to repellents were reduced in comparison to wild type flies. Our data also revealed that *HhTAR1* is more expressed in the male antennae of *H. halys* than in female ones. These results suggest that TAR1, besides being associated with olfactory repellence processes, could also play a role in responses to olfactory-reproductive stimuli, such as pheromones, or in mating behaviors (Mazzoni et al., 2017). The *HhTAR1* mRNA resulted more abundant in the two segments of flagellum with a 6-fold difference in comparison to the other antennal structures. A typical insect antenna contains numerous sensilla, essential structures for smell, taste, mechanoreception and thermo-hygro perception (Zacharuk, 1985). The great number of sensilla in the apical parts of the *H. halys* antennae correlates with the high *HhTAR1* expression level in the same areas, further strengthening a role for TAR1 in olfaction. Since the physiological role of each sensilla may be predicted based on their morphology, size and distribution (Keil, 1999), the *H. halys* sensilla were investigated by SEM. Different types of sensilla have been classified in the Pentatomidae, including basiconic, trichoid, coeloconic and chaetic sensilla (Brèzot et al., 1997). The most abundant structures in FL1 and FL2 segments of the adult *H. halys* were trichoid sensilla (TR) followed basiconic sensilla (BS) or “grooved sensilla” as observed also by Ibrahim et al. (2019) in the same insect. Both TR and BS-C share olfactory functions (Toyama et al., 2006) as suggested by the presence, on the surface, of distinctive microperforations necessary to connect the odorous molecules with the olfactory receptors in the OSNs (Zacharuk, 1985). It is difficult to associate each type of sensillum to a specific olfactory-mediated behavior, but the removal of both FLs completely inhibited the adult *H. halys* aggregation, indicating that these structures, and probably also TR and BS are necessary to perceive the aggregation pheromone (Toyama et al., 2006). On the other hand, sporadic BS have been observed in SC and both PEs along with structures identified as pit sensilla or coeloconic sensilla that could be involved in the thermo-hygrosensory reception (Altner & Prillinger, 1980). It is interesting to note that *HhTAR1* is more expressed in the FLs as compared to the SC and both Pes suggesting a correlation between TAR1 and olfactory sensilla. These data would therefore lead to hypothesize an important role for HhTAR1 in olfactory processes. Interestingly, *HhTAR1* showed high expression levels also in eggs and in 1^st^ and 2^nd^ instar nymphs, followed by a dramatic decrease from the 3^rd^ instar nymphs onwards. The results also revealed that between 2^nd^ and 3^rd^ instar nymphs the *HhTAR1* expression decreased more (about 80 %) in the abdomen and thorax tissues in comparison to the head. Also in adults, *HhTAR1* levels remained high in brain and antennae in comparison to other tissues. Previous studies observed that *H. halys* nymphs exhibit 4-times higher mortality than adults after treatment with essential oils for 1 or 48 h (Bergmann & Raupp, 2014) The high *HhTAR1* expression in CNS and antennae nymphs might explain the greater sensitivity to volatile compounds with insecticides properties, such as essential oils. In fact, TAR1 is a putative target for biopesticides, such as monoterpenes (Gross et al., 2017, Finetti et al., 2020). It is not yet clear how monoterpenes exert their toxicity in vertebrates, but their volatile nature is currently used to repel various insect pests (Reis et al., 2016).

The analysis on *HhTAR1* expression patterns together with the SEM observations on *H. halys* antennae strongly suggest a connection between HhTAR1 and *H. halys* olfactory regulation. To better investigate this aspect, *HhTAR1* was silenced by RNAi in young nymphs. In recent years, several Hemiptera genes have been successfully silenced through this method (Christiaens & Smagghe, 2014; Bansal et al., 2016; Ghosh et al., 2017; Lu et al., 2017; Mogilicherla et al., 2018, Riga et al., 2019). In these studies, the dsRNA was delivered exclusively by microinjection or by feeding but both these delivery methods are problematic. Microinjection requires experience and specific instruments to control the injected volume, as well as minimizing the wound that often causes a drastic increasing in mortality (Christiaens et al., 2020). In fact, through the microinjection we were able to obtain a *HhTAR1* RNAi downregulation in *H. halys* 2^nd^ instar nymphs (data not shown) but with an extremely high mortality. On the other hand, the dsRNA delivery by feeding requires a large amount of dsRNA and it does not allow to control the amount of dsRNA ingested by each insect (Joga et al., 2016). The dsRNA topical delivery has been recently tested in two Hemiptera species, *Diaphorina citri* and *Acyrthosiphon pisum*. In *D. citri*, 20 ng of dsRNA solution topically delivered on the abdomen were able to silence several *Cyp* genes by about 70-90% (Killiny et al., 2014). In *A. pisum*, 120 ng of dsRNA solution induced a downregulation of a target gene by 90 % after 24-36 hours (Niu et al., 2019). Accordingly, in *H. halys* 2^nd^ instar nymphs (rich in *HhTAR1* mRNA), a 100 ng dose of *HhTAR1* dsRNA topically delivered appeared sufficient to silence *HhTAR1* by about 50 % after 24 hours, as verified by RT-qPCR. The different RNAi efficiency observed between *D. citri, A. pisum* and *H. halys* could be explained to the different body structure: the abdominal cuticle of *H. halys* nymphs is thicker than that of *D. citri* and *A. pisum*, an aspect that could limit absorption of dsRNA solution. At any rate, this is the first time that RNAi mediated gene silencing is induced by topic delivery in *H. halys*. In previous studies, RNAi silencing has already been successfully performed on this insect using microinjection and feeding as delivery methods. One µg of dsRNA injected in *H. halys* adults was able to silence several target genes by 60-80 % after 72 h (Mogilicherla et al., 2018). On the contrary, when the dsRNA solution was delivered by feeding to *H. halys* 2^nd^ and 4^th^ instar nymphs, some target genes were silenced only by 40 - 80 % (Kumar et al., 2017). Although the dsRNA topically delivered results less efficient as gene silencer in *H. halys*, the administered amount of dsRNA is lower compared to the microinjection and the feeding applications. Reducing the dsRNA amount could be an effective strategy to prevent off target effects (Romeis & Widmer, 2020).

Upon *HhTAR1* silencing, *H. halys* 2^nd^ instar nymphs were tested in their olfactive performances by an innovative behavioral assay. This assay measured the repellent effect of (*E*)-2-decenal, one of the main alarm compounds released by *H. halys* under threats, on 2^nd^ instar nymphs (Zhong et al., 2017; Zhong et al., 2018; Nixon et al., 2018). The *HhTAR1*-dsRNA treatment caused a reduced sensitivity to (*E*)-2-decenal in comparison to the *LacZ*-dsRNA control nymphs. suggesting that the (*E*)-2-decenal-mediated alarm requires a functional TAR1.

In conclusion, HhTAR1 could play a relevant role in the *H. halys* olfactory network, contributing to modulate olfaction-mediated behaviors, such as reception of alarm pheromone compounds. A more detailed characterization of the interconnections between TAR1 and the olfactory system will pave the way for developing TAR1-targeting volatile compounds, such as essential oils, with both repellent and insecticidal properties against *H. halys*.

## Acknowledgement

The authors would like to thank Dott. Santolo Francati (University of Bologna) for providing *H. halys* adults, Dott.ssa Morena De Bastiani (University of Ferrara) for the technical assistance, Prof. Stefano Della Longa (University of L’Aquila) and Prof. Alessandro Arcovito (Università Cattolica del Sacro Cuore, Rome) for molecular modelling advice, Dott.ssa Federica Albanese (University of Ferrara) and Dott.ssa Milvia Chicca (University of Ferrara) for language revision.

This study was funded by ‘Camera di Commercio Industria, Artigianato e Agricoltura’ of Ferrara ‘Bando 2018’ and by the Emilia Romagna region within the Rural Development Plan 2014-2020 Op. 16.1.01 – GO EIP-Agri - FA 4B, Pr. “ALIEN.STOP”, coordinated by CRPV.

---

**Supplementary figure 1 (S1).**
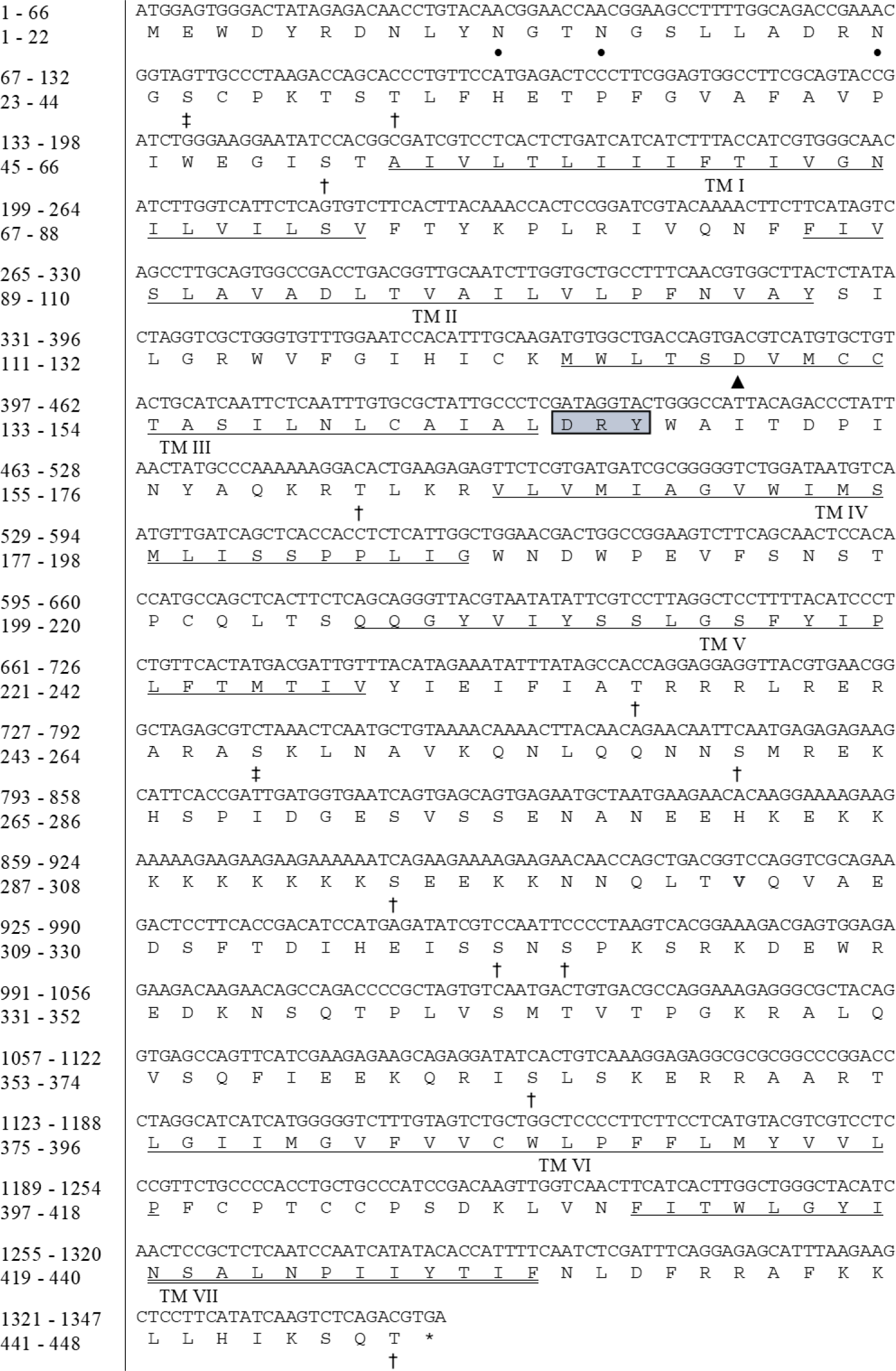
Nucleotide sequence of the TAR1 open reading frame cloned from *Halyomorpha halys* and deduced amino acid sequence. Prediction of the transmembrane segments (underlined and numbered from I to VII) was obtained with TMHMM v. 2.0 software. After the third transmembrane domain there is the DRY motif (highlighted with a box). Potential sites for N-linked glycosylation (predicted with NetNGlyc 1.0 server) are shown with a • and potential sites for PKA or PKC phosphorylation (predicted with NetPhos 3.1 server) are shown with a † and a ‡ respectively.

**Supplementary figure 2 (S2).**
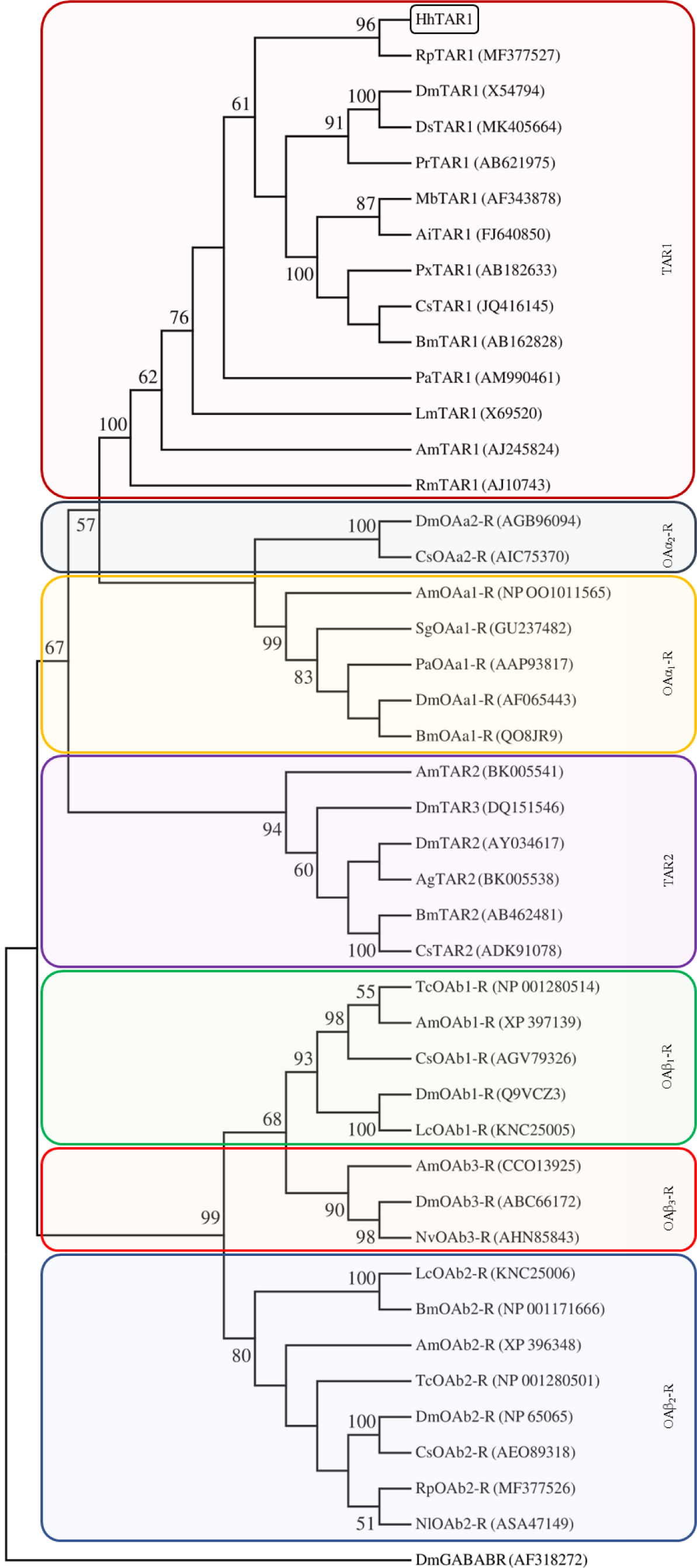
Phylogenetic relationships of HhTAR1 and other insect amine receptors resulting from neighbour joining analysis, using MEGA7. The values shown at the nodes of the branches are the percentage bootstrap support (1000 replications) for each branch. Alignment was performed using the amino acid sequences found in GenBank (accession number are indicated). *Drosophila melanogaster* GABA-B receptor (DmGABABR) was chosen as outgroup. Dm, *Drosophila melanogaster*; Ds, *Drosophila suzukii*; Pr, *Phormia regina*; Hh, *Halyomorpha halys*; Rp, *Rhodnius prolixus*; Px, *Papilio xuthus*; Cs, *Chilo suppressalis*; Bm, *Bombyx mori*; Ai, *Agnotis ipsilon*; Mb, *Mamestra brassicae*; Pa, *Periplaneta americana*; Lm, *Locusta migratoria*; Am, *Apis mellifera*; Rm, *Rhipicephalus microplus*; Sg, *Schistocerca gregaria*; Ag, *Anopheles gambiae*; Tc, *Tribolium castaneum*; Nv, *Nilaparvata lugens*; Lc, *Lucilia cuprina*; Nl, *Nilaparvata lugens*.

**Supplementary figure 3 (S3).**
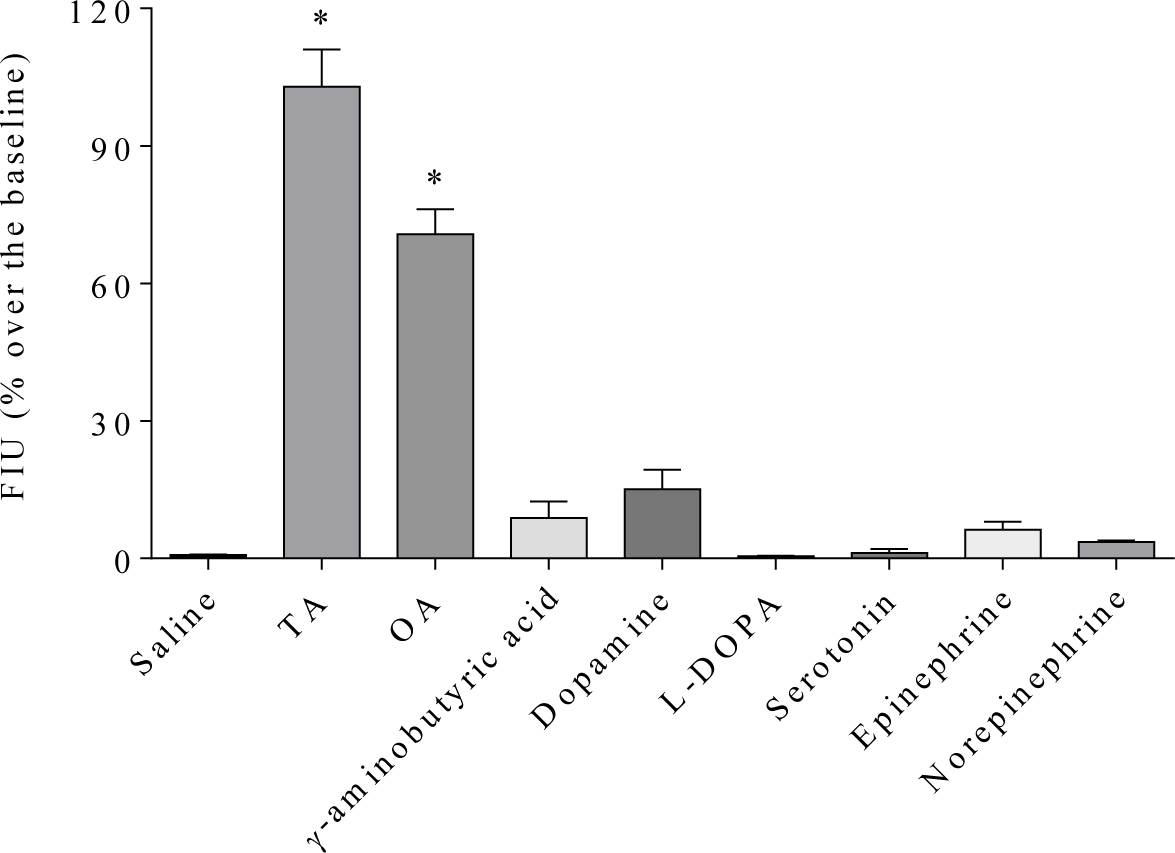
Effect of biogenic amines and γ-aminobutyric acid on the intracellular calcium release in HEK 293 stably expressing HhTAR1. All compounds were tested at 10^−4^ M. Data represent means ± SEM of three separate experiments performed in duplicate. * p < 0.001 vs saline according to one-way ANOVA followed by Dunnett’s multiple comparison test.

**Supplementary figure 4 (S4).**
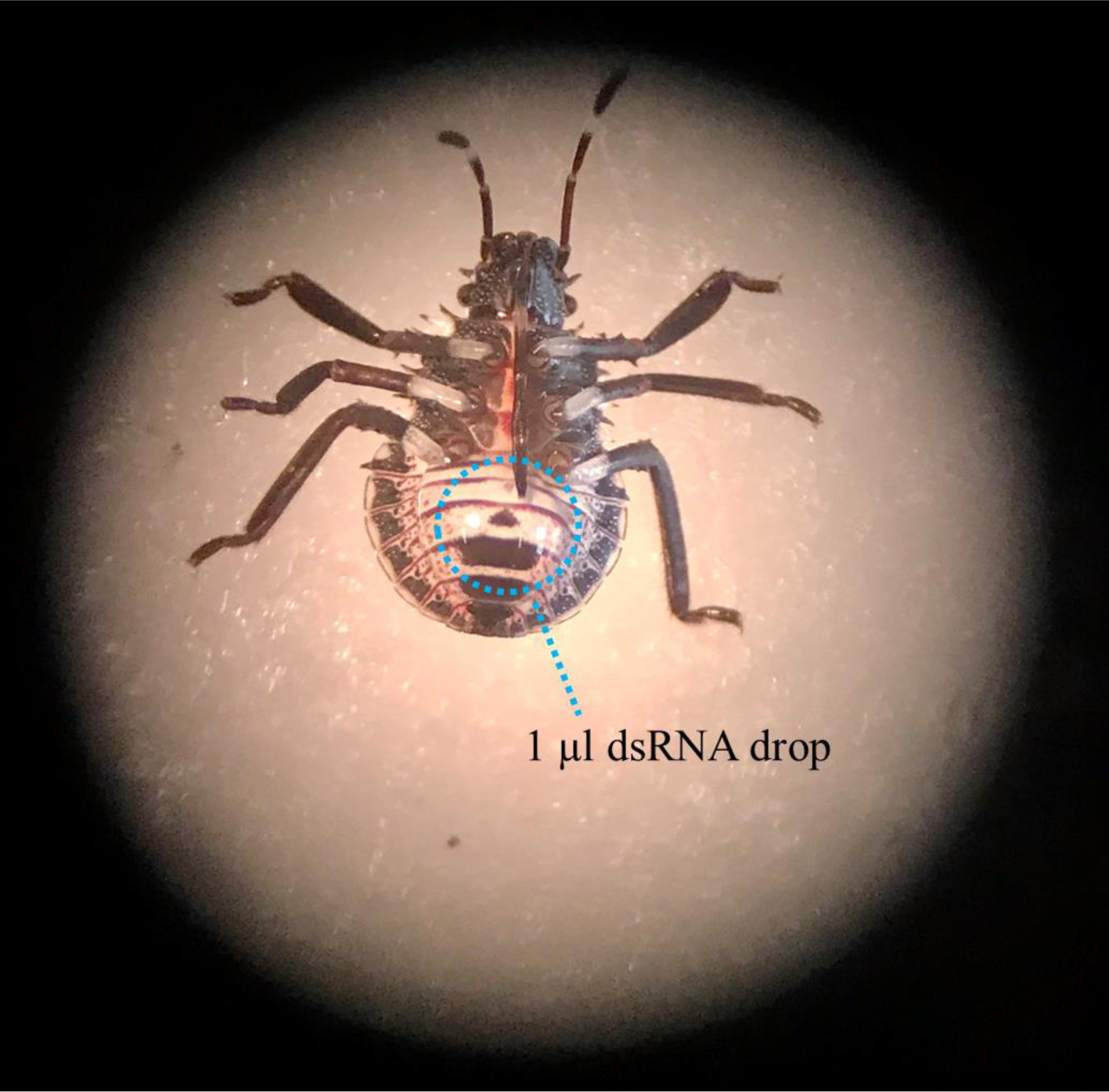
Image of dsRNA topically delivered on a *H. halys* 2^nd^ instar nymph. The 2^nd^ instar nymphs were collected 3 days post-ecdysis and placed on double-sided adhesive tape to avoid movements. One µl of the dsRNA solution was placed on the dorsal side of the abdomen. When the dsRNA solution was completely absorbed, the nymphs were put back in the nursery cage.

